# crsra: A package for Cleaning and Analyzing Coursera Research Export Data

**DOI:** 10.1101/275537

**Authors:** Aboozar Hadavand, Jeffrey Leek

## Abstract

Due to the fundamental differences between traditional education and Massive Open Online Courses (MOOCs) and the ever-increasing popularity of MOOCs more research is needed to understand current and future trends in education. Although research in the field has rapidly grown in recent years, one of the main challenges facing researchers remains to be the complexity and messiness of the data. Therefore, it is imperative to provide tools that pave the way for more research on the new subject of MOOCs. This paper introduces a package called **crsra** based on the statistical software R to help clean and analyze massive loads of data provided by Coursera. The advantages of the package are as follows: a) faster loading and organizing data for analysis, b) an efficient method for combining data from multiple courses and even across institutions, and c) provision of a set of functions for analyzing student behaviors.

## Introduction

Research on Massive Open Online Courses (MOOCs) is young. Bozkurt et al. (2017) studied literature published on MOOCs through 2015 and found that the number of articles published on the subject increased from 1 in 2008 to 170 in 2015. More research is needed to fully understand the effectiveness, reach, limits, and the potential of MOOCs. However, the main challenges in studying MOOCs remains to be data. Data on MOOCs are not usually publicly available since it is owned by private providers and there are concerns about the privacy of students. More importantly, as Lopez et al. (2017) point out, the size and complexity of MOOC data is an overwhelming challenge to many researchers. Therefore, it is imperative to provide tools that pave the way for more research on the new subject of MOOCs.

This paper introduces a package called **crsra** (Hadavand and Leek, 2018) based on the statistical software R to help clean and analyze massive loads of data from the Coursera MOOCs. Coursera is one of the leading providers of MOOCs and was launched in January 2012. With over 25 million learners, Coursera is the most popular provider in the world being followed by EdX, the MOOC provider that was a result of a collaboration between Harvard University and MIT, with over 10 million users. Coursera has over 150 university partners from 29 countries and offers a total of 2000+ courses from computer science to philosophy (Coursera, 2018). Besides, Coursera offers 180+ specialization, Coursera’s credential system, and four fully online Masters degrees. A typical course on Coursera includes recorded video lectures, graded assignment, quizzes, and discussion forums.

Since the early years of the company, Coursera has encouraged researchers to analyze students’ data and has facilitated the use of the data and the platform for A/B testing. Starting November 2015 Coursera introduced a dashboard for self-service data exports. Through this tool, partner institutions and instructors could download data for a single course or all courses associated with the institution. Research data exports are sets of CSV files and are designed for use in relational database systems. One of the advantages of the data is the existence of a single *hashed user ID* for each student. This user ID is consistent for learners across all courses offered by an individual institution and allows for connecting learner grades and progress across courses.

The advantages of the package are as follows: a) faster loading of data for analysis, b) efficient method for combining data from multiple courses and even across institutions,^1^ and c) provision of a set of functions for analyzing student behaviors.

### Coursera Research Data

There are five types of research data export for each course. Table 1 summarizes these five types. The data are written in roughly 100 tables containing information such as course content, students’ demographic data, progress, and outcomes, and forum data. Figure 1 shows different types of research data exports provided by Coursera.

**Table 1:** Types of research data export

| Data Type | Description |
| --- | --- |
| Assessment submission data | Assessment submissions of quizzes, peer review, and programming assignments by learners. |
| Course grade data | Contains the highest grade achieved by each learner on each required assessment as well as the time stamp of the learner’s highest-scoring submission. This table also includes each learner’s overall grade in the course. |
| Course progress data | Contains data documenting the time stamp for when the learner interacted with each piece of course content and the time stamps for when items were opened, completed, reopened, reattempted, etc. |
| Demographic data | Contains the following information for all enrolled learners: general geographical data (based on IP address), browser language preference, and information for learners who completed their learner profile responses or participated in Coursera’s platform-wide demographic survey (including age, gender, education level, and employment status). |
| Discussion data | Contains forum activity data such as posts, responses, upvotes/downvotes, flags, and questions and answers associated with course content items. |

**Figure 1:**
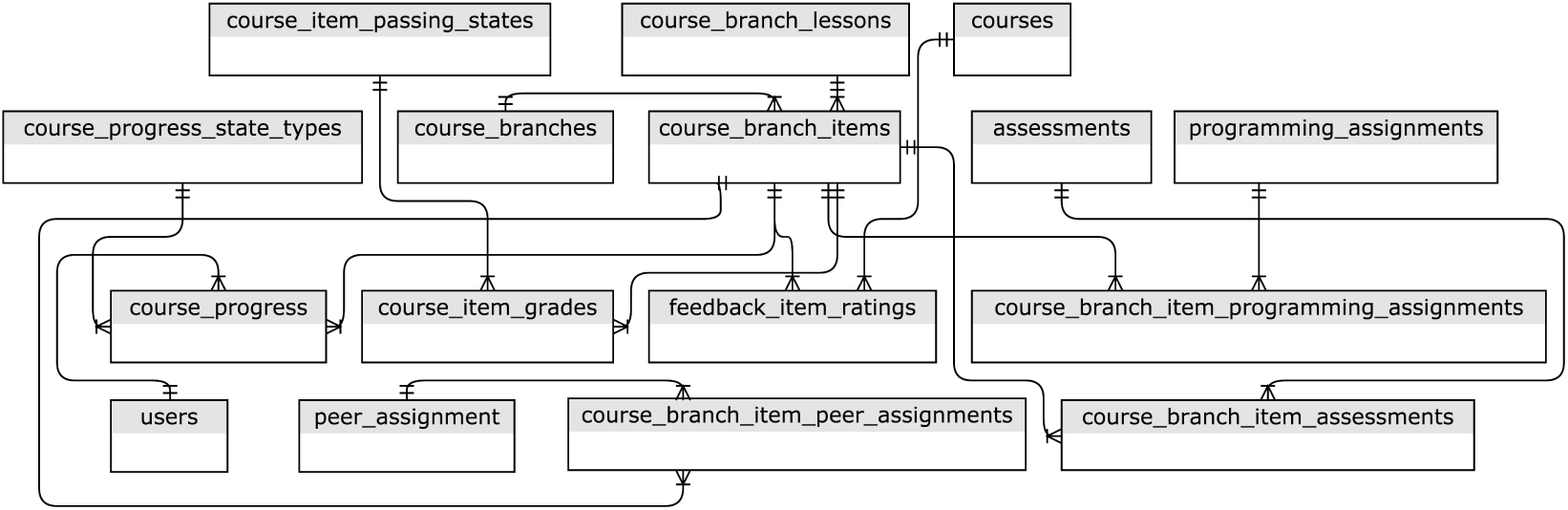
The major relationships between table groups, with minor connections omitted (Source: Coursera)

While Coursera provides tools for creating PostgreSQL databases in a docker container,^2^ as we mentioned earlier, importing data for analysis remains to be a challenge for researchers with limited experience with relational databases. Moreover, such tools are usually not platform independent.^3^

#### The crsra Package

The **crsra** package helps import and organize Coursera’s research data exports into R. It also runs some preliminary analysis on the data. In the following section, we introduce the package and provide instruction on how to import Coursera research data exports. To install this package, you will need to install **devtools** (Wickham et al., 2017), then execute the following commands to install the **crsra** package.

~~~
library(“devtools”)
devtools::install_github(“jhudsl/crsra", build_vignettes = TRUE)
~~~

To import your data dump into R, first, point your working directory to the directory that contains all the unzipped course folders. Then execute the command crsra_import().^4^ If you are not pointing to the correct directory, you will receive a warning, and the execution will be halted. Note that the data import may take some time if the course data is large and there are several courses in your working directory. Also note that by running the crsra_import() command, you import all tables for each course into R in a list called all_tables.^5^

Tables can be called using all_tables `[["course_name"]][["table_name"]]`. For instance, if you like to call the `table peer_comments` in the course Regression Models, you can simply execute all_tables`[["Regression Models"]][["peer_comments"]]`. To see a list of courses imported by the `crsra_import()` command check the variable coursenames. To see a list of all the tables check the variable tablenames.

To see the data import in use, we use the package on data from Johns Hopkins University (JHU) Data Science Specialization on Coursera. This specialization, developed by Jeffrey Leek, Roger Peng, and Brian Caffo, consists of ten courses. There have been more than two million enrollments since the launch of this program in April 2014. The size of data on the students who took these ten courses since 2015 is around 18 gigabytes. In the following example, we use the **crsra** package to import a Coursera data dump at our disposal on all the courses and to find the number of students who passed a specific course item (course item 67c1O) in the course “Regression Models.”

~~~
library(dplyr)
all_tables %>%
.[["Regression Models"]] %>%
.[["course_item_grades"]] %>%
dplyr::filter(course_item_id == “67c1O”) %>%
dplyr::filter(course_item_passing_state_id == 2) %>%
dplyr::summarise(n = n())
# A tibble: 1 x 1
# n
# <int>
# 1 8640
~~~

The package also includes a few other functions in addition to the main crsra_import() function. A list of functions and their descriptions is provided in Table 2.

**Table 2:** Some of the functions in the **crsra** package

| Function | Description |
| --- | --- |
| `crsra_membershares` | Returns a summary of the total number and the shares of users in each course broken down by factors such as roles, country, language, gender, employment status, education level, and student status. |
| `crsra_gradesummary` | Returns total grade summary broken down by the factors mentioned above. |
| `crsra_progress` | Summarizes, for each course item, the total number and the share of users who ceased activity at that specific course item. The function ranks course items by their attrition. |
| `crsra_assessmentskips` | Users may "skip" reviewing a peer-assessed submission if there is a problem with it. This function categorizes skips by their type such as "inappropriate content", "plagiarism", etc. The function also returns a word cloud appeared in peer comments as to why they skipped the submission. |
| `crsra_timetofinish` | Calculates the time to finish a course for each user. |

We can also use the function `crsra_gradesummary()` to calculate the average student grade for the courses in the data import. By using the argument groupby we can calculate average grades for different learner subgroups based on gender, education, student status, employment status, and country. For instance, the following analysis returns the average overall course grade for male and female learners in the course *The Data Scientist’s Toolbox*. The results show that female learners’ grades are on average 6 points lower on a 100 scale than male learners’ grades.

~~~
crsra_gradesummary(groupby = “gender”) %>%
.[["The Data Scientist’s Toolbox"]]
#Note that maximum grade possible is 1.
# A tibble: 2 x 2
# reported_or_inferred_gender AvgGrade
# <chr> <dbl>
#1 male 0.7250660
#2 female 0.6691554
~~~

#### A Preliminary Analysis of Student Behavior on Coursera

The existence of fundamental differences between traditional education and MOOCs has attracted a new wave of studies on students’ behavior and outcomes in the online world. These differences are best reflected in the definition of MOOCs by McAuley et al. (2010) that “[a]n online course with the option of free and open registration, a publicly shared curriculum, and open-ended outcomes which integrates social networking, accessible online resources … and most significantly builds on the engagement of learners who self-organize their participation according to learning goals, prior knowledge and skills, and common interests.” Such differences require further research on MOOCs. Understanding how students progress through an education program is critical for any educational planning and decision making (King, 1972). Models of student progress are needed to estimate the probability of a student completing a particular item in a course and predict the time required to finish a course. Furthermore, conventional measures of academic success and progress cannot be defined in the same way for MOOCs. For instance, as Perna et al. (2014) states, we have limited knowledge on whether learners’ progress through a MOOC should be measured in a sequential fashion or in a way that captures the flexibility and freedom in learning behavior that is unique to MOOCs.

There are only a handful of studies on students’ progress and outcomes in MOOCs. Perna et al. (2014) perform a descriptive analysis of student progress through a set of 16 courses on Coursera. They found that most users accessed course content in the sequential order defined by the instructor of the course. Ho et al. (2014) study 17 courses taught on EdX and found that most of the attrition in online courses happen in the first week of course activity (about 50 percent attrition) and that the average percentage of learners who cease activity in the second week declines sharply to 16 percent. Most of these studies are specific to a set of courses or platforms. Due to the many differences in the characteristics of MOOCs, any extrapolation of the results to MOOCs in general has to be done with caution.

In the following section, we will investigate students’ progress through the ten Data Science Specialization courses on Coursera provided by JHU. Using the crsra_timetofinish function in the **crsra** package, we can first investigate the time difference between the first and last activities in a course for each student. Time to finish is only calculated for those who completed the course. Figure 2 depicts the density plot for time to complete for three of the courses in the specialization. Note that the density plot varies across courses. While for *Developing Data Products* and *Getting and Cleaning Data* a majority of students finish the courses in around 30 days, for *Data Science Capstone* a majority of students finish the course in 50 days.

~~~
TTF <- crsra_timetofinish()
~~~

**Figure 2:**
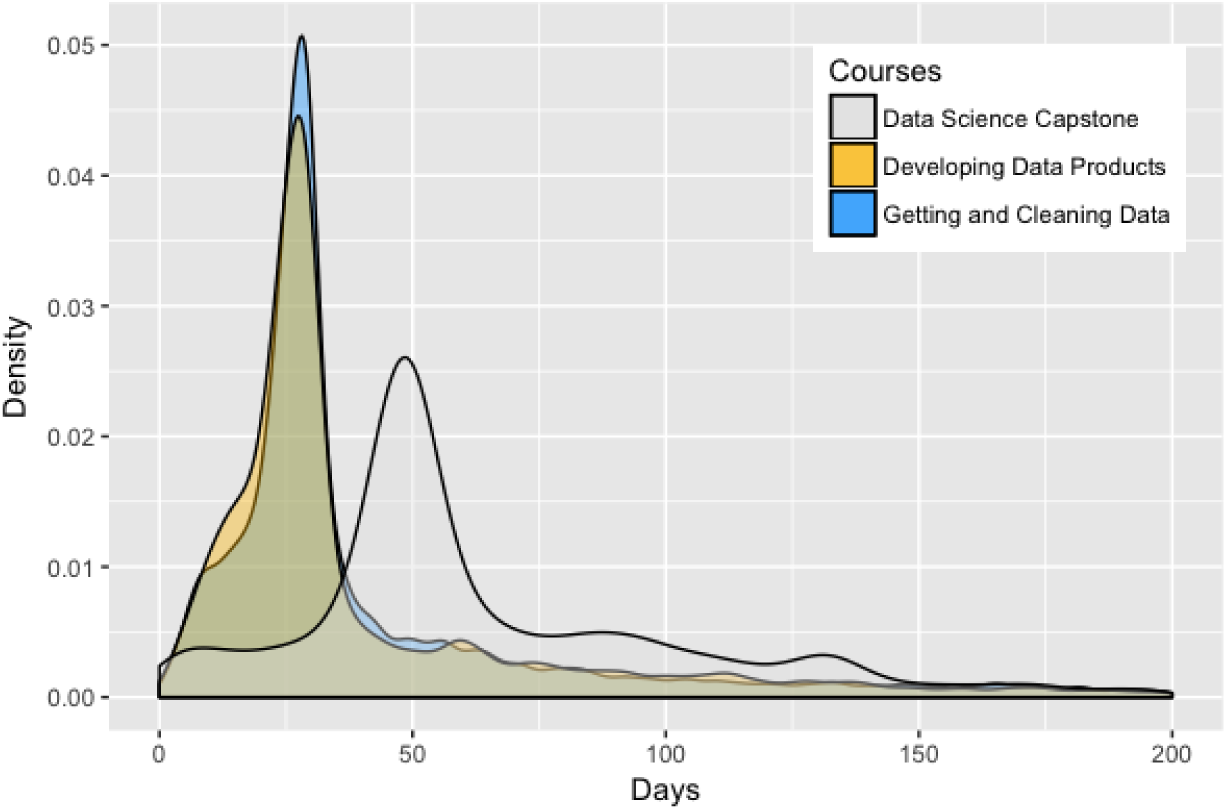
Density plots for time to finish defined as the time difference between the first and last activities across three courses.

In the table called users, Coursera provides a field for student status of the learner including full-time and part-time students and those who are not degree students. We can look at how time to finish is different for groups with different student status. Figure 3 reports this for the course *Getting and Cleaning Data* and shows that part-time students take longer to finish the course.

~~~
TTF.Status <- TTF %>%
.[["Getting and Cleaning Data"]] %>%
dplyr::left_join(all_tables[["Getting and Cleaning Data"]][["users"]],
by = “jhu_user_id", ‘copy’=TRUE) %>%
dplyr::filter(!is.na(student_status))
~~~

**Figure 3:**
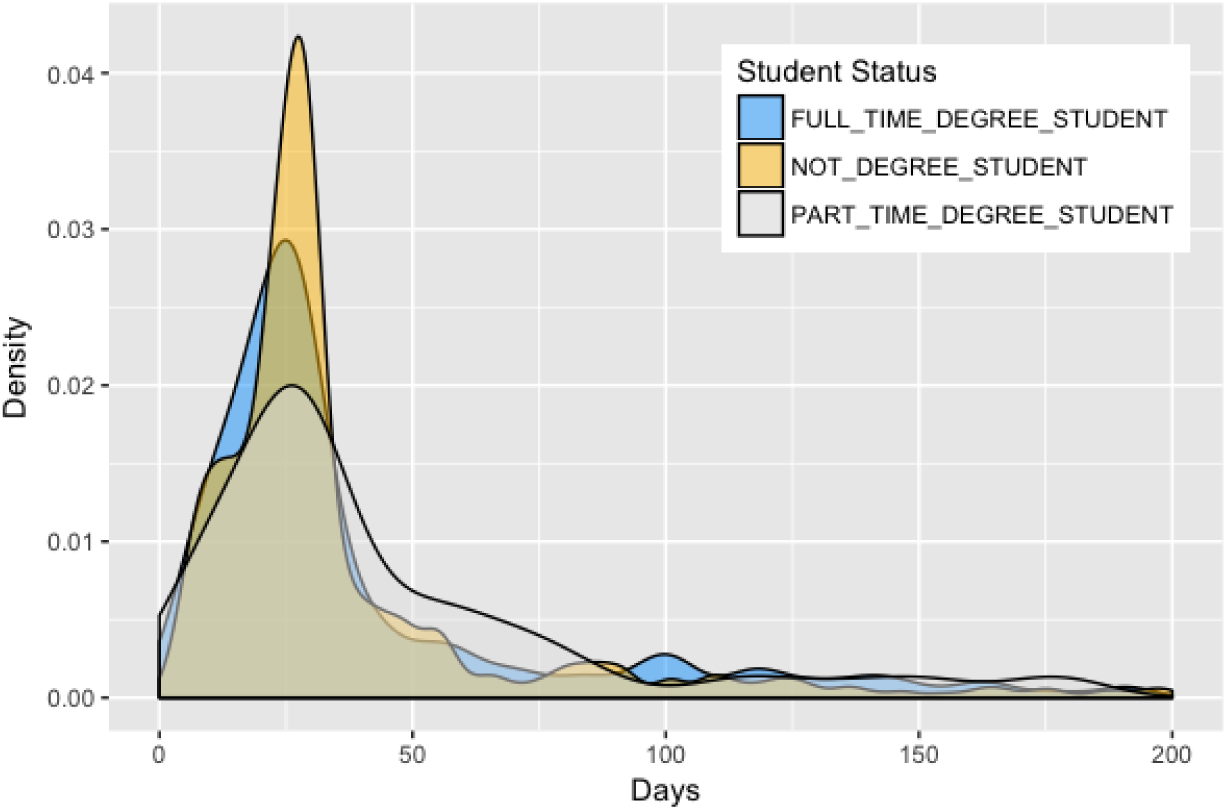
Density plots for time to finish for learners with different student statuses in the course Getting and Cleaning Data.

Our next step is to understand student progress in the courses. One of the factors that distinguish MOOCs from traditional classrooms is the flexibility in advancing through the course. While in traditional education the class length, pace, and completion dates are determined by the instructor of the course, in MOOCs it is the student who, for the most part, has the freedom to choose these factors. We can then look at how many course items students pass in the first week of course activity. One obvious but yet interesting finding is large variations across students. For instance, if we look at the course *Getting and Cleaning Data*, we can use the following code to find the number of course items completed in the first week of course activity. The course has roughly 40 items including lectures and assignments. The variable nweek1 captures the number of passed course items in the first week of course activity, calculated as one week after a student’s first activity in the class. The density plot in Figure 4 represents the variations across students. For a majority of learners, the number of passed course items in the first week is two. However, the number of those who finish more than ten items in the first course is significant. Also interesting is the double-peak shape of the density plot. It is interesting to see that there are more people who stop after completing twelve course items in the first week than there are who stop after completing seven course items. This indicates an interesting structural change in students’ pace between course items 7 and 12.

~~~
passed.items <- all_tables %>%
.[["Getting and Cleaning Data"]] %>%
.[["course_progress"]] %>% dplyr::group_by(jhu_user_id) %>%
# 604800 is the number of seconds in a week
dplyr::filter(course_progress_ts <= min(course_progress_ts) + 604800) %>%
dplyr::summarise(nweek1 = n())
~~~

**Figure 4:**
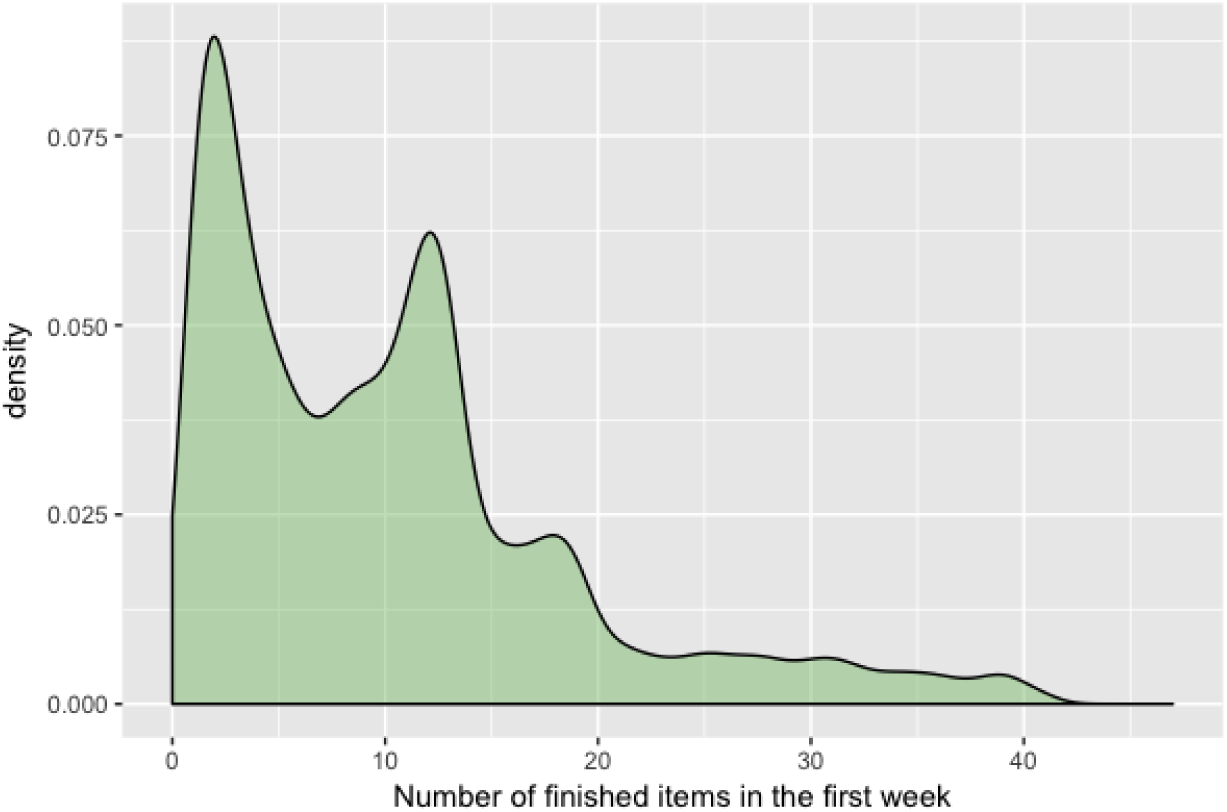
Density plot for the number of passed items in the first week of course activity for the course Getting and Cleaning Data.

A third interesting variable to look at when studying students’ progress in MOOCs is the time gaps between each session. In this exercise, we looked at the time lapsed between each two consecutive course items for each learner throughout the course *Getting and Cleaning Data*. We used the following code for the analysis. Note that we have ranked course items based on the time stamp when a student passes them and not based on their natural order defined by the course instructor.

~~~
gaps <- all_tables %>%
.[["Getting and Cleaning Data"]] %>%
.[["course_progress"]] %>%
# 2 is an indicator that the course item is completed
dplyr::filter(course_progress_state_type_id == 2) %>%
dplyr::group_by(jhu_user_id, course_item_id) %>%
# This is for keeping only the latest event for each course item
dplyr::filter(course_progress_ts == max(course_progress_ts)) %>%
dplyr::ungroup() %>%
dplyr::arrange(jhu_user_id, course_progress_ts) %>%
dplyr::group_by(jhu_user_id) %>%
# This is for converting the time gap to hours
dplyr::mutate(time.dif = as.numeric(course_progress_ts - lag(course_progress_ts))/3600) %>%
dplyr::filter(!is.na(time.dif)) %>%
dplyr::filter(time.dif != Inf | time.dif != -Inf)
~~~

Figure 5 shows student progress in the course Getting and Cleaning Data for three sample students. The vertical axis is the gap between two consecutive sessions in hours. These three students are chosen intentionally to show three different learning paths. Panel A shows progress for a student with short gaps between sessions for the first half of the course and longer gaps towards the end. We call this pattern “slowing down” pattern. This pattern is typical of many students in this course. Panel B shows progress for a student with short gaps between sessions in the beginning and the end of the course and longer gaps in the middle. Students in this group are not as common as the first group. Finally, Panel C shows progress for a student with no apparent pattern in their progress throughout the course. Only a small group of students follow this pattern in our data.

**Figure 5:**
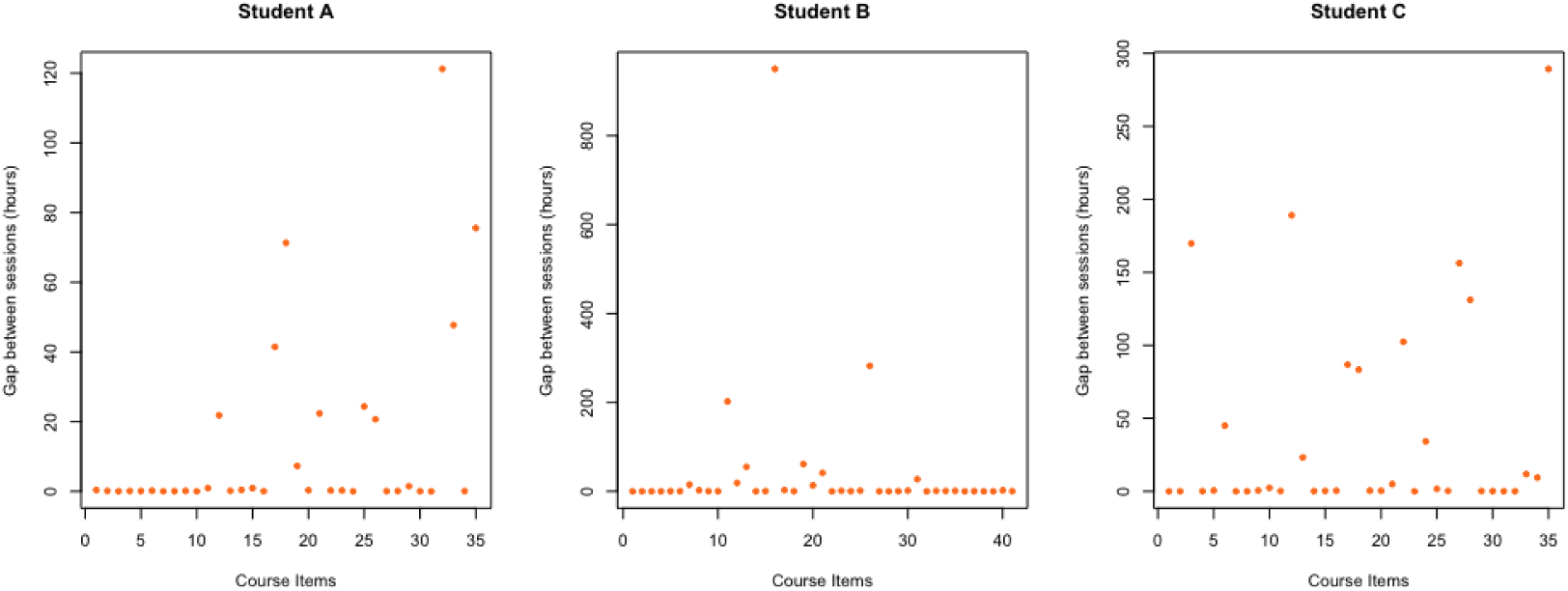
Student progress through the course Getting and Cleaning Data for three sample students. The vertical axis shows the time gap between completing an item and the next item in hours.

Let us look at the average gap between sessions for the first and the second half of the course. We can then calculate how much the average session gap changes from the first half to the second. Across our sample of students who registered for the course, the average change in session gap from the second half to the second half is positive and equal to 132 percent. In other words, the gap between session more than doubles from the first half of the course to the second half. Figure 6 shows the density plot of this statistic across our sample. The long right tail in the Figure supports the fact that most students follow the slowing-down pattern. However, the Figure also shows that there are students who speed up during the second half of the semester.

**Figure 6:**
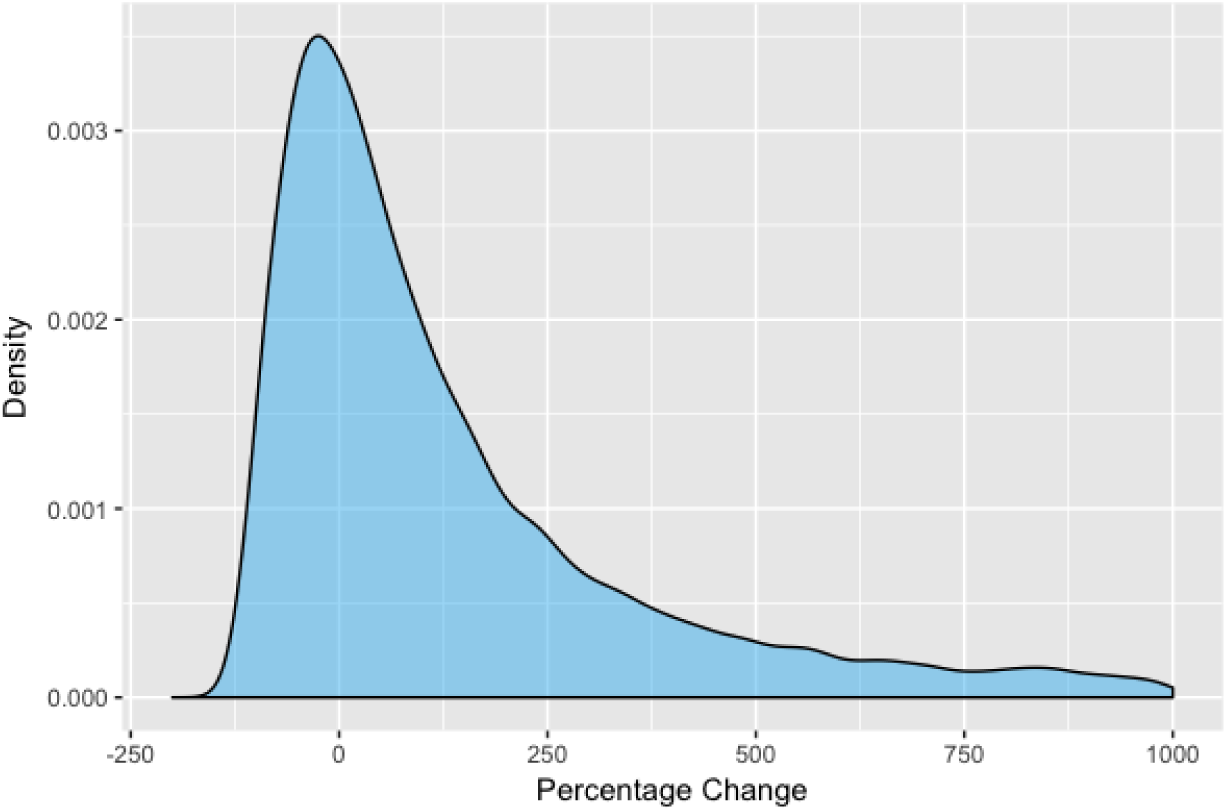
Percentage change in time gaps in course progress between the first and the second half of the course Getting and Cleaning Data.

We can do a similar analysis for some subgroups of our sample. Some of the most interesting categories are gender, educational attainment, and whether the learner paid for taking the course. The variable was_payment in the table users_courses certificate_payments captures whether the learner has ever paid for a course certificate. This purchase could be a “single payment” for the course or a “bulk payment” for a specialization that contains the course. The following code is used for the analysis of payers and non-payers and the results for all categories are shown in Table 3.

~~~
gaps.payment <- gaps %>% dplyr::group_by(jhu_user_id) %>%
dplyr::summarise(avgtime = mean(time.dif)) %>%
dplyr::inner_join(all_tables[["Getting and Cleaning Data"]][["course_grades"]],
by = “jhu_user_id", ‘copy’=TRUE) %>%
dplyr::filter(course_passing_state_id %in% c(1, 2)) %>%
dplyr::left_join(
all_tables[["Getting and Cleaning Data"]][["users_courses certificate_payments"]],
by = “jhu_user_id", ‘copy’=TRUE) %>%
dplyr::filter(!is.na(was_payment)) %>%
dplyr::group_by(was_payment) %>%
dplyr::summarise(avggap=mean(avgtime))
~~~

**Table 3:** Gaps between sessions for different subgroups of learners in the course Getting and Cleaning Data Course

| Categories | Average gap in hours |
| --- | --- |
| Gender | Female: 39<br>Male: 36 |
| Educational Attainment | Less than high school: 152<br>High school diploma: 39<br>College (no degree): 39<br>Associate degree: 23<br>Bachelor's degree: 32<br>Master's degree: 38<br>Professional degree: 22<br>Doctoral degree: 34 |
| Paid for the course? | Yes: 36<br>No: 39 |

The last exercise in our analysis is to study how Coursera’s change in policy from a pay-per-course business model to a subscription model changed students’ progression throughout the course. On October 30, 2016, Coursera introduced a new payment system through which they allowed students to purchase access to all content in a specialization on a month-by-month or annual basis (Coursera, 2016). As a result, the student would only pay for the amount of time they need to learn the material. This system replaced the existing model where students would pay up front for each course regardless of how long it took them to finish the course. A question to ask is whether the switch to this system where payments are tied to the length of time it takes students to complete the class speeds up learning paces. The following code calculates the average number of course items passed in the first week of activity for the two groups: those who enrolled in the course before October 30, 2016, and those who joined after. We hypothesize that those who pay monthly are more likely to finish more items in the first week than those who pay a fixed price.

~~~
passed.items.policy <- passed.items %>%
dplyr::left_join(all_tables[["Getting and Cleaning Data"]][["course_memberships"]],
by = “jhu_user_id", ‘copy’=TRUE) %>%
dplyr::filter(!is.na(course_membership_ts)) %>%
dplyr::mutate(subscription = ifelse(course_membership_ts < “2016-11-01 00:00:00",
“before", “after”)) %>%
dplyr::group_by(subscription) %>%
dplyr::summarise(subnw = mean(nweek1))
~~~

The results suggest that those who enrolled before the policy change on average passed three courses less than the group who enrolled after (9 versus 12). Figure 7 shows the density plot of the number of passed items in the first week of activity for the two groups. This comparison, however, has a caveat: there is some selection bias since those who enrolled before, and those who registered after October 2016 may be fundamentally different.

**Figure 7:**
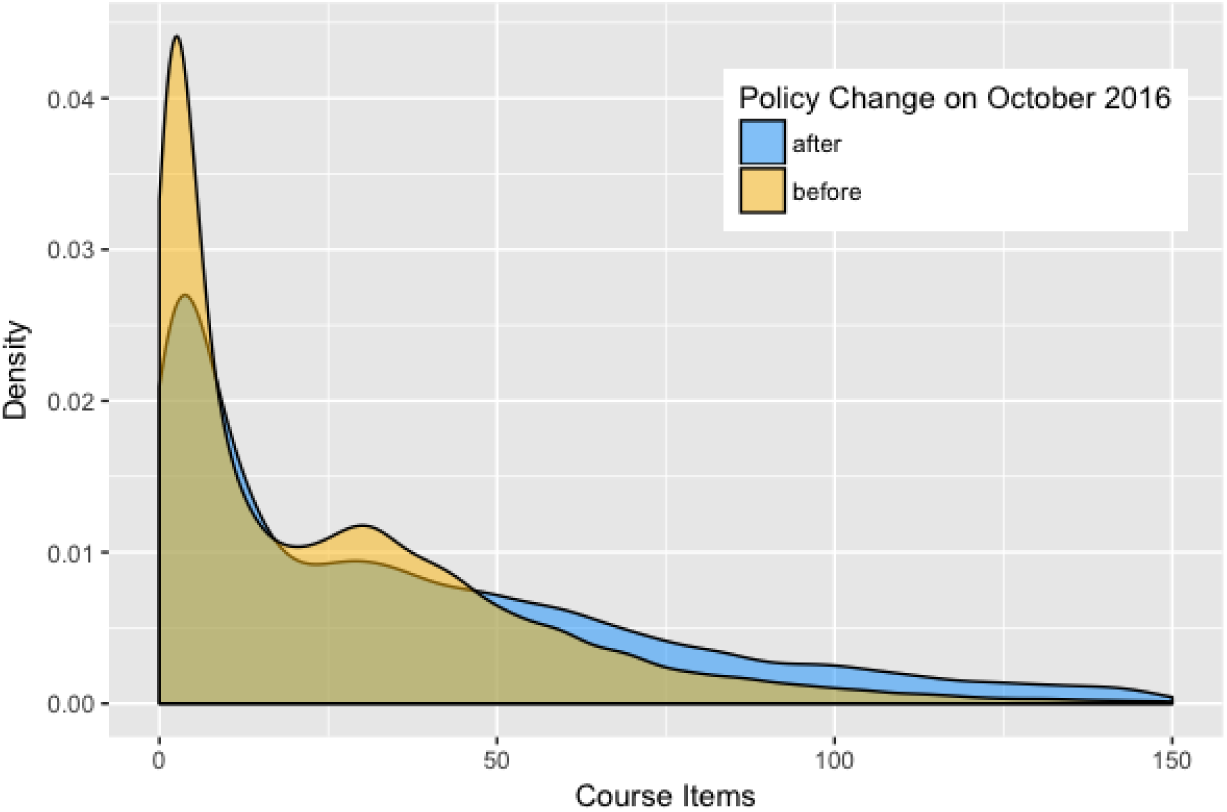
Density plots for the number of passed items in the first week of course activity for Getting and Cleaning Data for those who enrolled in the course before October 30, 2016, and those who enrolled after.

## Discussion

The **crsra** package provides tools for cleaning and analysis of Coursera’s research data exports. Our motivation for the development of the package was initially to analyze student progress using JHU Data Science Specialization data. The size and messiness of data were our main challenges. In this regard, we felt the need for a tool that imports the data into R, cleans the data, and provides some analysis of the courses. We, later on, decided to make the tool available to all researchers who use Coursera’s research data.

Because MOOCs are new, analyzing students’ behaviors on MOOCs are essential. One of the main differences between MOOCs and traditional college classrooms is how students progress through MOOCs. While the order and length of course content and the pace at which they are taught are chosen by the instructor in classrooms, in MOOCs, it is the student who determines when and how to learn the material. We, therefore, used the **crsra** package to analyze student progress and pace in one of a sample of courses offered by JHU. We hope the package and the analysis provided in this paper will pave the way for future studies on student behaviors on MOOCs.

## Footnotes

1 This is important since although MOOC researchers have access to thousands of students in their sample, few studies benefit from data across multiple courses and institutions. Such analysis helps draw more robust conclusions about student behaviors (Reich, 2015).

2 The tool is called ‘courseraresearchexports’ and can be found at https://github.com/coursera/courseraresearchexports

3 In an initial version of **crsra** based on PostgreSQL, we had the problem of some team members not being able to set up the database correctly on their PCs.

4 You can also dowanload the dummy data included in the data folder in the package repository on Github.

5 For a list of all the tables in the data download, please go to the URL https://github.com/jhudsl/crsra/blob/master/ListofTables.md

